# Bio-informed Protein Sequence Generation for Multi-class Virus Mutation Prediction

**DOI:** 10.1101/2020.06.11.146167

**Authors:** Yuyang Wang, Prakarsh Yadav, Rishikesh Magar, Amir Barati Farimani

**Affiliations:** Department of Mechanical Engineering, Carnegie Mellon University, Pittsburgh, PA 15213 USA; Department of Biomedical Engineering, Carnegie Mellon University, Pittsburgh, PA 15213 USA; Machine Learning Department, Carnegie Mellon University, Pittsburgh, PA 15213 USA

## Abstract

Viral pandemics are emerging as a serious global threat to public health, like the recent outbreak of COVID-19. Viruses, especially those belonging to a large family of +ssRNA viruses, have a high possibility of mutating by inserting, deleting, or substituting one or multiple genome segments. It is of great importance for human health worldwide to predict the possible virus mutations, which can effectively avoid the potential second outbreak. In this work, we develop a GAN-based multi-class protein sequence generative model, named ProteinSeqGAN. Given the viral species, the generator is modeled on RNNs to predict the corresponding antigen epitope sequences synthesized by viral genomes. Additionally, a Graphical Protein Autoencoder (GProAE) built upon VAE is proposed to featurize proteins bioinformatically. GProAE, as a multi-class discriminator, also learns to evaluate the goodness of protein sequences and predict the corresponding viral species. Further experiments show that our ProteinSeqGAN model can generate valid antigen protein sequences from both bioinformatics and statistics perspectives, which can be promising predictions of virus mutations.

## 1 Introduction

Viral diseases have been a threat to public health worldwide, like the current outbreak of the Severe Acute Respiratory Syndrome Coronavirus (SARS-CoV-2) [1, 2], which has already caused a loss of hundred thousands of human lives and more than 6 million people infected worldwide [3]. One of the reasons that viral diseases are hard to prevent and control is the high mutability of viral genomes, especially for those belonging to the positive-sense single-stranded ribonucleic acid (+ssRNA) viruses family. +ssRNA viruses are more likely to mutate by inserting, deleting, or substituting one or multiple RNA segments because they lack stable double-stranded deoxyribonucleic acid (DNA) structures [4]. By mutation, viruses can become more lethal and more infectious, which poses an even greater threat to healthcare. Therefore, effective and accurate prediction of potential virus mutations can play a vital role in developing effective therapeutic antibodies against the virus and preventing a viral infection from becoming a pandemic [5].

One common way to identify viruses is through the unique proteins synthesized by their genomes. The protein is composed of 20 standard amino acids that appeared in the genetic code, and each can be represented by a single letter via FASTA format coding [6]. A commonly targeted viral protein is the spike protein, which acts as the antigen, and the antibody binds to this antigen with very high specificity [7, 8, 9], as shown in Fig. 1(a). This process of highly selective interactions between the antigen and antibody forms the basis of antibody-mediated virus neutralization [10]. The antigen, therefore, can be the unique identity for the virus, which is closely related to the immune reaction [11, 12]. Additionally, antigen change can also be an efficient and effective prediction of the corresponding virus genome mutation.

**Figure 1:**
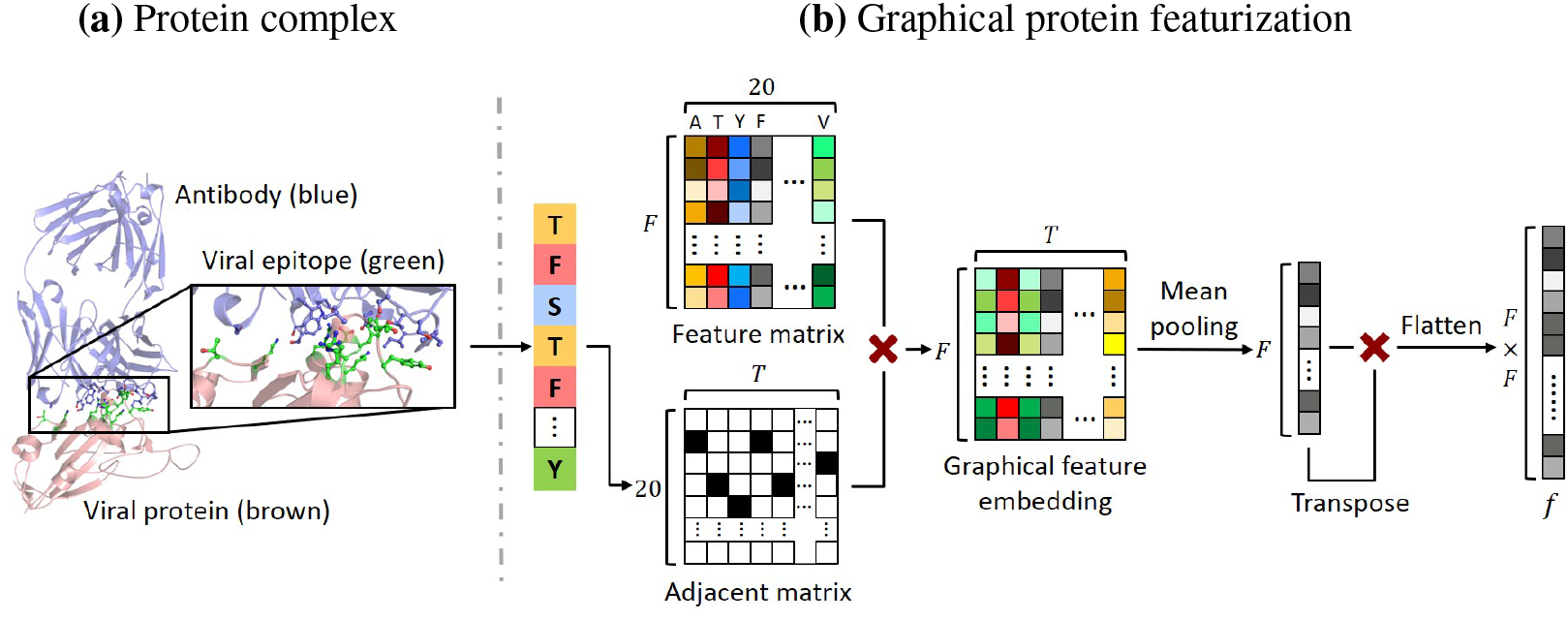
**(a)** The protein complex is the from the PDB ID 2DD8 (SARS-CoV virus spike protein and neutralizing antibody) [18]. The blue protein at top is the antibody molecule and the brown molecule at bottom is the viral protein. The amino acids in green color are the viral epitope amino acids which are involved in recognition by the antibody, are shown in the zoomed-in sub figure. **(b)** Graphical featurization of FASTA coding of the epitope, where the nodes refer to the amino acids and edges are the bonds in between.

In this work, we propose ProteinSeqGAN, a multi-class protein sequence generative model based on Generative Adversarial Networks (GANs) [13], which contains the generator, *G*, and the Graphical Protein Autoencoder (GProAE). The sequence generator, conditioned on viral species, is built upon Recurrent Neural Networks (RNNs) [14] and takes in random noise input to predict the antigen epitope synthesized by the viral genomes. The GProAE treats the input protein sequence as an undirected graph [15, 16] and featurizes it to a 1444-dimensional representation. The representation is then fed into a Variational Autoencoder (VAE) [17] to encode and reconstruct the input. Through the parameter-sharing encoder, the representation is also leveraged to evaluate the goodness of protein sequences and classify it into viral categories, which are both backpropagated to train the multi-class generator. Experiments from both bioinformatics and statistics perspective prove that our ProteinSeqGAN model can generate valid amino acid sequences, which can be reliable viral mutation predictions corresponding to different viral species. Such predictions are of great importance in curing viral diseases and preventing potential second outbreaks.

## 2 Related Works

### 2.1 Generative Adversarial Network

Since first proposed, Generative Adversarial Networks (GANs) [13] have been a trend in generating realistic samples through the min-max game between a discriminator and a generator. Conditional GAN (cGAN) is then developed to generate samples conditioned on discrete labels [19, 20], images [21, 22] and texts [23, 24]. To improve the performance of conditional generation cross multiple domains, StarGAN [25] proposes to add a domain classification besides the single validity in the discriminator. Wasserstein GAN (WGAN) [26] and WGAN-GP [27] provides an alternate way of training the generator model to better approximate the distribution of data observed in a given training dataset. Besides, Variational Autoender (VAE) [17] is leveraged to improve generative performance of GAN [28, 29].

GAN has also been introduced to sequence generation. In [30], the WGAN [26] and WGAN-GP [27] is combined with Recurrent Neural Networks (RNNs) [14], which learns to generate realistic sentences. To bypass the differentiation problem in the generator with gradient descent, SeqGAN [31] models the generator as a stochastic policy in reinforcement learning (RL). StepGAN [32] pushes even further by not only evaluating the full generated sequence but also evaluating sub-sequences at each generation step to provide more RL signals. Besides RL, the feature matching between the high-dimensional latent features from real and generated sequences can also be leveraged to improve sequence generation as introduced in TextGAN [33].

### 2.2 Protein Generation and Validation

Proteins are large biomolecules composed of one or more chains of amino acids, as shown in Fig. 1(a). Protein structure analysis and methodical novel protein design are of great importance in understanding molecular and cellular biological functions. However, the generation of protein sequences is complicated by the vast number of combinations. For a protein of length *T*, there are a total number of 20^*T*^ possible amino acid sequences, which are infeasible to exhaust. Conventionally protein sequences were generated by wet-lab experimentation, which involved cell culture and proteomic assays to validate the synthesized protein [34]. Such methods are time-consuming and expensive due to the wet-lab experimentation and fail to leverage the available data and computational approaches to accelerate such experimentation [35, 36].

Recently, GANs have been introduced to generate novel proteins. In [37], GAN is leveraged to generate protein structures for fast *de novo* protein design, which encodes protein structures with pairwise distances between *α*-carbons on the protein backbone. Further, in [38], cGAN is utilized in protein generation given graph representations of 3D structure. GAN has also been implemented on protein design in novel folds [39], where protein is generated conditioned on low-dimensional fold representations. Besides, [40] leveraged GAN generated proteins as a data augmentation to improve protein solubility prediction results.

Additionally, validation of proteins is a central problem faced in bioinformatics [41, 42]. Multiple methods have been developed to validate the protein sequences [43, 44], such as BLOSUM matrix (BLOcks SUBstitution Matrix) [45] which validates the amino acid sequences by quantitating the probability of substitution mutations. In [46], a statistical method is proposed, in which a natural vector is developed for amino acid sequences to evaluate the validity.

## 3 Method

In this section, we introduce ProteinSeqGAN, a multi-class protein sequence generative model. The problem is that given the viral species *c* of virus families 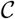, developing a generative model, which takes random noise *z* and viral species *c* as input to generate the sequence of amino acids *A* = {*a*_1_,…, *a_t_*,…, *a_T_*} of length *T*, where 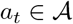, and 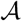 is the single-letter FASTA format [6] of all the 20 standard amino acids along with the end of sequence token. The generated protein sequences are closely related to the viral genomes, which can be essential for virus mutation prediction.

### 3.1 Graphical Protein Featurization Autoencoder

To incorporate the bioinformatics of proteins, a Graphical Protein Featurization Autoencoder (GProAE). GProAE treats the protein sequence as an undirected graph, where the nodes represent amino acids in the protein, and the edges are the bonds formed between them. As shown in Fig. 1(b), given the input sequence of length *T*, an adjacent matrix of dimension (20, *T*) is built, where each column is the one-hot encoding of the amino acid. The feature matrix contains *F* features of all the 20 standard amino acids, which is of dimension (*F*, 20). Features selected to describe each amino acid, include hydrophilicity, aromaticity, orientation of side chains, number of hydrogen donors, number of carbon atoms, *etc*., which span a wide range of amino acid properties and captures variance between different amino acids. A detailed description of all the 38 bioinformatics features is provided in Appendix A.1. The graphical feature embedding is then computed by the matrix multiplication of the adjacent matrix and feature matrix. Subsequently, mean pooling operation is conducted on each row of the graphical feature embedding, which outputs a homogeneous (*F*, 1) vector for sequences with variant lengths. The pooled vector is again multiplied with the transpose of itself to generate a (*F, F*) matrix. At the last step, the matrix is flattened to get the graphical feature *f*.

After we get the feature *f* through the graphical protein featurization block, an autoencoder is built to learn the latent features *z* of proteins with an encoder-decoder architecture, as shown in Fig. 2(b). The encoder is first developed with a parameter-sharing embedding block, which maps *f* to a lower-dimensional vector *e* = *Emb*(*f*). The embedding e is then fed into an encoder to predict the mean *μ* and variance *σ* of the Gaussian distribution, and the latent vector *x* is sampled from the Gaussian distribution as 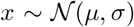. *x* is then fed into the decoder to reconstruct the graphical feature through 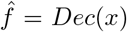. Additionally, to train a GAN model, validity of multi-class protein sequences is required. Therefore, the embedding e is streamed down to two discriminators, *D_val_* and *D_cls_*. The former validates the goodness of input protein sequences by computing *D_val_*(*e*), while the latter works as a classifier to predict the viral species corresponding to the input through *D_cls_*(*e*). Such classification helps to evaluate the generated proteins conditioned on various viral species.

**Figure 2:**
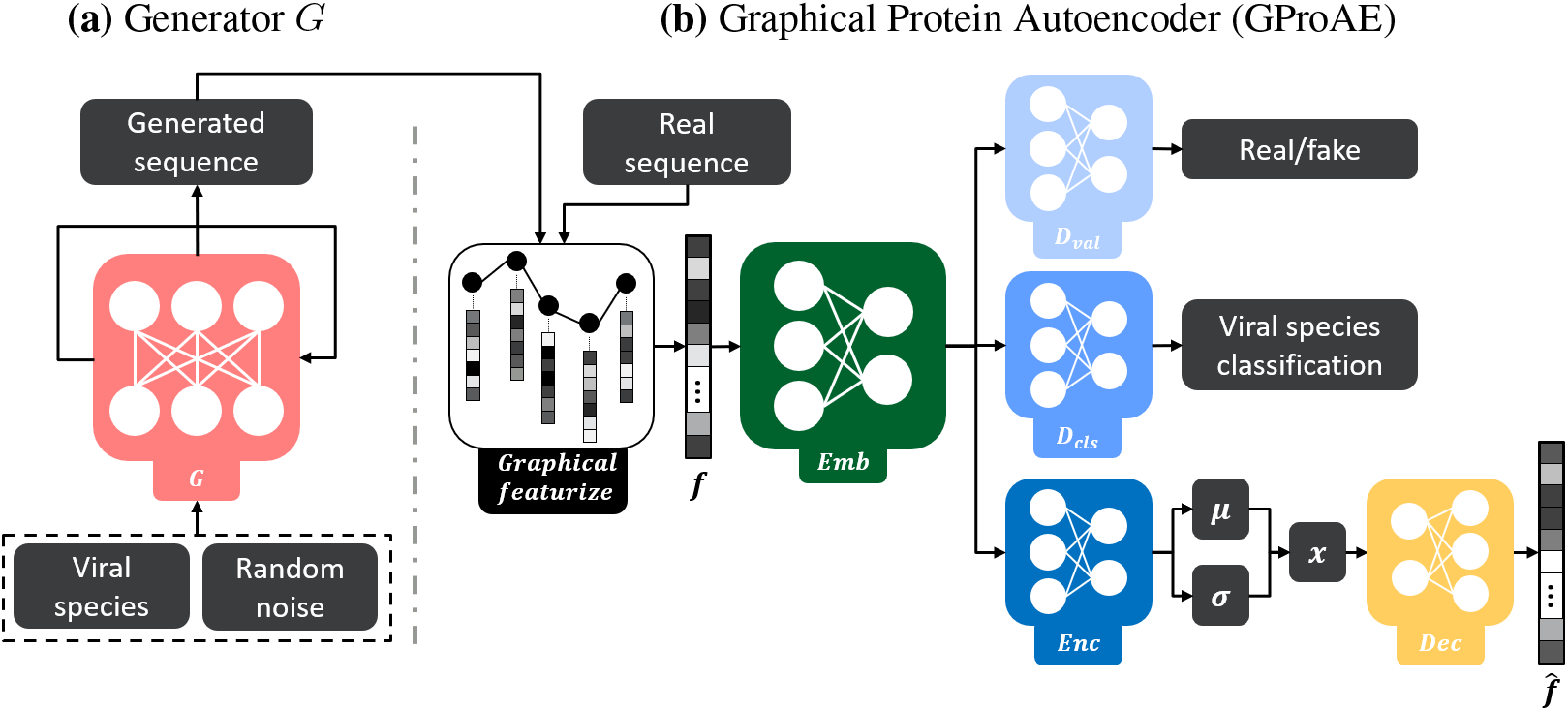
Overview of ProteinSeqGAN model: **(a)** The generator *G*, built upon RNN, takes in as input both the viral species c and a random noise *z* to generate the protein sequence. **(b)** GProAE graphically featurizes the input protein to *f*, and learns the representation through the parametersharing embedding block. *D_val_* and *D_cls_* predicts the goodness and viral category respectively, while the encoder and decoder compute the reconstructed graphical feature 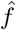.

### 3.2 Conditional Protein Sequence Generator

Recurrent neural networks (RNNs) [14] are used to model the conditional sequence generator, *G*. As illustrated in Fig. 2(a), *G* takes the embedding of the virus species as the intial hidden state *h*_0_. Also a random noise *z*, following an isotropic Gaussian distribution 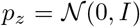, is fed into the RNN model as the input at the initial step. Let *x_t_* denote the input to the RNN model at position *t*, the hidden state *h_t_* is recursively calculated by the update function *g*:

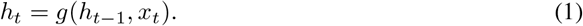

Afterwards, a softmax function and the nonlinear ouput function *o* maps the hidden state *h_t_* to the probability distribution of all the 20 amino acids and the end of sequence token at position *a_t_*:

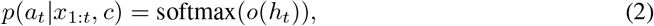

where *x*_1_ = *z*, and *x_t_* = *p*(*a*_*t*-1_|*x*_1:*t*-1_, *c*) for *t* > 1, which is the output amino acids distribution at the previous position *t* – 1.

During training, the model generates the amino acid sequence by selecting the token with the highest probability at each position as given in Eq. 2, and truncating the sequence at the first appearance of the end of sequence token. While during test, the amino acids are sampled from the output distribution *p*(*a_t_*|*x*_1:*t*_, *c*). In our implementation, LSTM [14] is used to model the update function *g* to alleviate the gradient vanishing and exploding problems, which are common in RNN training.

### 3.3 Full Model

Shown in Fig. 2 is the whole pipeline of ProteinSeqGAN model, which is composed of the conditional sequence generator, *G*, and the multi-class discriminator, GProAE. GProAE takes as input both sequences from the training set and the generated sequences from *G*, and featurizes the proteins graphically. The *D_val_* and *D_cls_* then evaluates the quality and classifies the viral species respectively. While *G* tries to generate realistic protein sequences by cheating the *D_val_* and *D_cls_* in GProAE. To this end, we introduce the objective functions utilized to train the two components jointly.

#### Adversarial Loss

To generate realistic protein sequences which are indistinguishable from real ones, adversarial loss is implemented based on WGAN [26]:

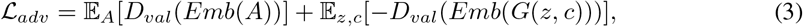

where *G* generates a protein sequence *G*(*z, c*) from a random noise *z* conditioned on viral species *c*. The validity score of proteins is computed via the embedding block *Emb*(·) and the discriminator *D_val_*(·). The GProAE is trained to maximize the adversarial loss, while the sequence generator G tries to fool the discriminator by minimizing the objective.

#### Protein Classification Loss

To better evaluate the conditional generation performance, an auxiliary classifier *D_cls_* [25] is developed in GProAE. The objective built upon cross-entropy function contains two components, a classification loss 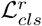 for real sequence input to train the GProAE, and a classification loss 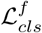 for generated fake sequence input to train *G*. The 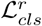 is defined as:

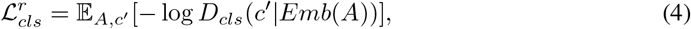

where the pair (*A, c′*) is the real protein sequence and its corresponding viral species from the training dataset. *D_cls_*(*c′*|*Emb*(*A*)) measures the predicted probability distribution over the actual viral category *c′*. By minimizing the objective 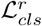, *D_cls_* learns to correctly classify the proteins, and meanwhile the parameter-sharing embedding block is also updated to obtain informative representations. Similarly, the classification objective for fake input 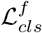 is defined as:

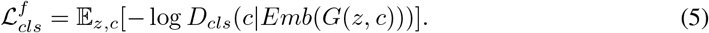

By minimizing 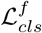, *G* tries to fool *D_cls_* by classifying the conditional generated sequences into the corresponding category *c*.

#### Feature Reconstruction Loss

An encoder-decoder architecture [17] is also modeled to encode and reconstruct the graphical feature *f* in a self-supervised manner. Since protein sequences are different in length which can be challenging to be fed directly to the encoder-decoder architecture. In detail, the feature reconstruction loss measures the *ℓ*_1_ distance between the graphical feature *f* and the reconstructed 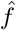 as given:

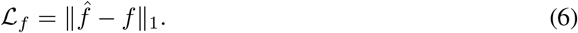

#### Prior Loss

To regularize the training process of GProAE, the learned latent feature *x* is assumed to follow an standard Gaussian distribution 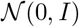. The prior loss computes the Kullback–Leibler (KL) divergence between the predicted Gaussian and the standard Gaussian as given:

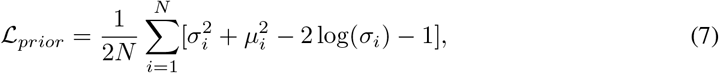

where *N* is the dimension of *x*, and *μ*, *σ*^2^ are the predicted mean and standard deviation respectively.

#### Sequence Reconstruction Loss

To accelerate and stabilize the training of *G*, we introduce the sequence reconstruction loss 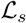. Namely, at position *t*, instead of the output from previous step *t* – 1, the amino acid from the a real sequence is fed into the LSTM cell, and *G* is supposed to reconstruct the amino acid sequence of length *T* correctly. Let 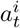 denote the one hot encoding of real amino acid at position *t*, where 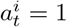 for 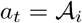 and 0 otherwise. Similarly, 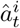 represents the predicted probability of amino acid type 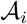 at position *t*. To evaluate the difference between the reconstruction sequence and the corresponding real sequence, a cross entropy loss is leveraged:

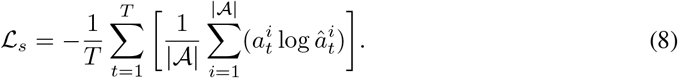

#### Total Objective

Overall, an affine combination of multiple loss functions are computed and backpropagated to update the model weights. The total objective to optimize the GProAE model in our settings is defined as:

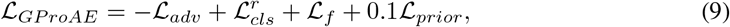

where the GProAE is trained by the losses 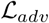 and 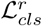 from GAN-based discriminators *D_val_* and *D_cls_* respectively, along with the losses 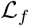 and 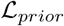 from the VAE model. While the objective for the generator *G* is given as:

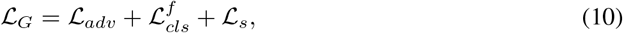

where *G* is trained to fool the discriminators by minimizing the adversarial loss 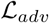 and the fake classification loss 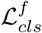, and meanwhile learns to reconstruct the protein sequence via 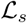.

## 4 Experiments

### 4.1 Dataset

The dataset to train out ProteinSeqGAN model is an amalgamation of the CATNAP dataset (Compile, Analyze, and Tally NAb Panels) [47], the VirusNet dataset [48] and the recent SARS-CoV-2 virus sequence [49], comprising of 1934 viral antigen sequences from over 16 different virus species. Viral antigen amino acids are selected within 5-7 angstroms from the antibody, which ensures that only the epitope of the antigen is selected, and mutations in this region can significantly alter the efficacy of antibodies. More details concerning the dataset can be found in the Appendix A.2.

### 4.2 Baselines

As our baseline models, we adopt the sequence generative model [30], which is implemented in combination of WGAN [26] and LSTM [14]. Also, we investigate the effects of different components in training our ProteinSeqGAN model.

#### WGAN & LSTM

Based on [30], both the discriminator and the generator are modeled directly upon an LSTM [14], where the discriminator evaluates the protein sequences and backpropagate that to the generator through the WGAN-based adversarial loss [13, 26].

#### ProteinSeqGAN without 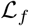

The encoder-decoder architecture [17] to reconstruct the graphical protein features *f* is deprecated. *f* is only fed to *D_val_* and *D_cls_*, which evaluates the goodness of the protein sequences and predicts the corresponding viral species respectively.

#### PorteinSeqGAN without 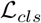

Viral classification discriminator *D_cls_* is removed from our ProteinSeqGAN in this setting. Namely, 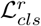 and 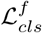 are both set to zero during training. The generator is trained only on the adversarial loss 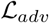 and sequence reconstruction loss 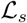.

### 4.3 Training

All models are trained using RMSProp [50, 51] with learning rate of 5 × 10^−5^ and *α* = 0.99. The generator is updated once after the GProAE updates ten times. All weights in the GProAE are clipped to [−0.01,0.01] after each update [26]. Batch size is set to be 16, and the total number of epochs is set to be 3000 in all the experiments. All LSTM hidden state and input noise prior dimensions are set to 64. In addition, the dimension of latent encoding vector *x* is set to 16.

### 4.4 Evaluation Metric

To ensure that the model has learned to generate antigen sequences which have a higher probability of being observed biologically, we used two independent sequence evaluation metrics, bioinformatics based validation, and statistics based validation. When used in tandem, these evaluation metrics ensure that the generated sequences are rigorously evaluated for their biological validity.

#### Bioinformatics Validation

The bioinformatics based evaluation begins with the sequence alignment of a generated viral antigen sequence with all the antigen sequences of that specific virus species in the dataset, which is based on Needleman–Wunsch algorithm [52]. The best matching alignment is then selected, which consists of the closest sequence in the dataset to the generated sequence. The substitution mutations between the best matching sequence from the dataset and the generated sequence are scored by the BLOSUM62 matrix [45], as shown in Fig. 3. To determine invalid sequences, sequence alignment of generated sequence and best match sequence is screened. If the generated sequence has a BLO-SUM invalid mutation, it is then labeled as an invalid sequence.

**Figure 3:**
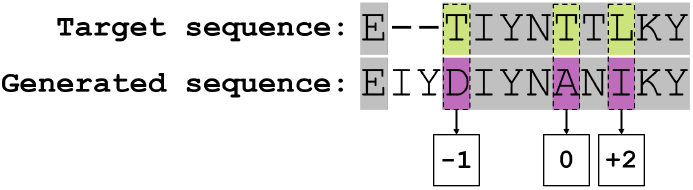
Bioinformatics validation via sequence alignment and BLOSUM scoring

#### Statistics Validation

Statistics based validation is based on the natural vector *v_N_* for each amino acid sequence [46]. *v_N_* is consists of three components: (1) the number of occurrence of the amino acid “k”, *n_k_*, (2) the mean distance of the amino acid “k” from the first position, *μ_k_*, and (3) the second normalized central moment of the distribution for the amino acid “k”, 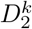. Therefore, the natural vector *v_N_* for an arbitrary amino acid sequence is given as:

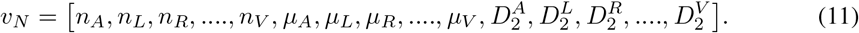

The validation criterion is based on a convex hull created by the bounds of *v_N_* from all the protein sequences in our dataset. Details and visualization of convex hull validation are provided in the Appendix A.3. These criteria ensure a strict check on the generated sequences and grants high-confidence in the biological validity of the sequences.

### 4.5 Generated Protein Sequences

The generated protein sequences and the corresponding viral species are illustrated in Table 1, where BV denotes the bioinformatics validity, and SV denotes the statistics validity. Also, we use T for true and F for false in the validity. Our ProteinSeqGAN can generate realistic protein sequence mutations comparing to the real proteins. The generated antigen sequences capture the pattern of different viral species like the approximate length and the common start amino acids. Also, there is no unrealistic fragment like the repeated appearance of the same amino acid in a row, which has been observed in other baseline models.

**Table 1:**
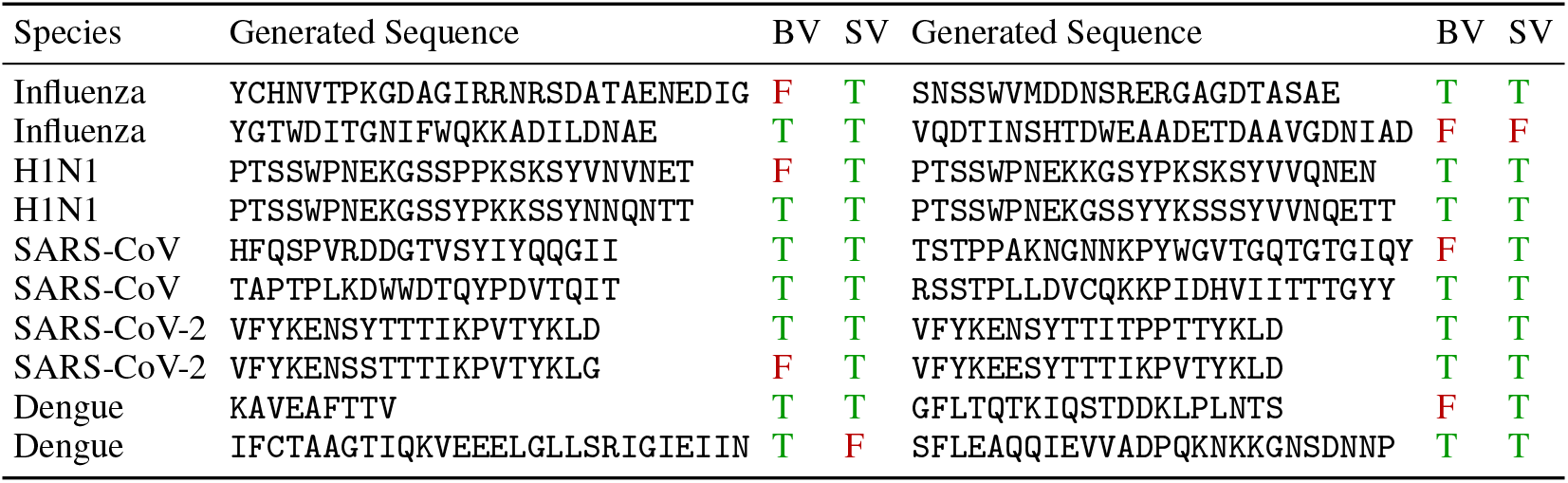
Generated protein sequences and validity evaluations

Table 2 shows the validity of all the 8,000 generated sequences for each model; namely, 500 antigen sequences are generated for each viral species. Among all the models, the full ProteinSeqGAN possesses the highest validity ratio from both bioinformatics (60.90%) and statistics validation (59.49%), along with the combination of these two criteria (36.10%). Without the GProAE architecture, WGAN + LSTM can barely generate realistic protein sequences, which indicates graphical featurization plays an essential role in evaluating the protein sequences where the LSTM discriminator fails to learn. Table 2 also demonstrates that the absence of 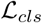 or 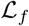 harms our generative model since the representations learned by the discriminators are not as informative as the full model.

**Table 2:** Validation for generated sequences from different models

| Model | BV (%) | SV (%) | Both valid (%) |
| --- | --- | --- | --- |
| WGAN + LSTM [30] | 22.17 | 25.77 | 6.62 |
| ProteinSeqGAN (without $\mathcal{L}_f$) | 48.86 | 64.80 | 39.04 |
| ProteinSeqGAN (without $\mathcal{L}_{cls}$) | 53.07 | 67.02 | 43.64 |
| ProteinSeqGAN | **60.90** | **77.61** | **47.00** |

In Fig. 4, we investigate the generated sequences in the graphical feature *f* domain, visualized using t-Distributed Stochastic Neighbor Embedding (t-SNE) [53]. Comparing to ProteinSeqGAN without 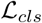, the full model generates sequences that are more closely related to the original viral species. Additionally, it is also illustrated the graphical features obtained in GProAE are representative in distinguishing antigen sequences from various species. Visualization of protein sequences from all the 16 viral species can be found in Appendix A.4.

**Figure 4:**
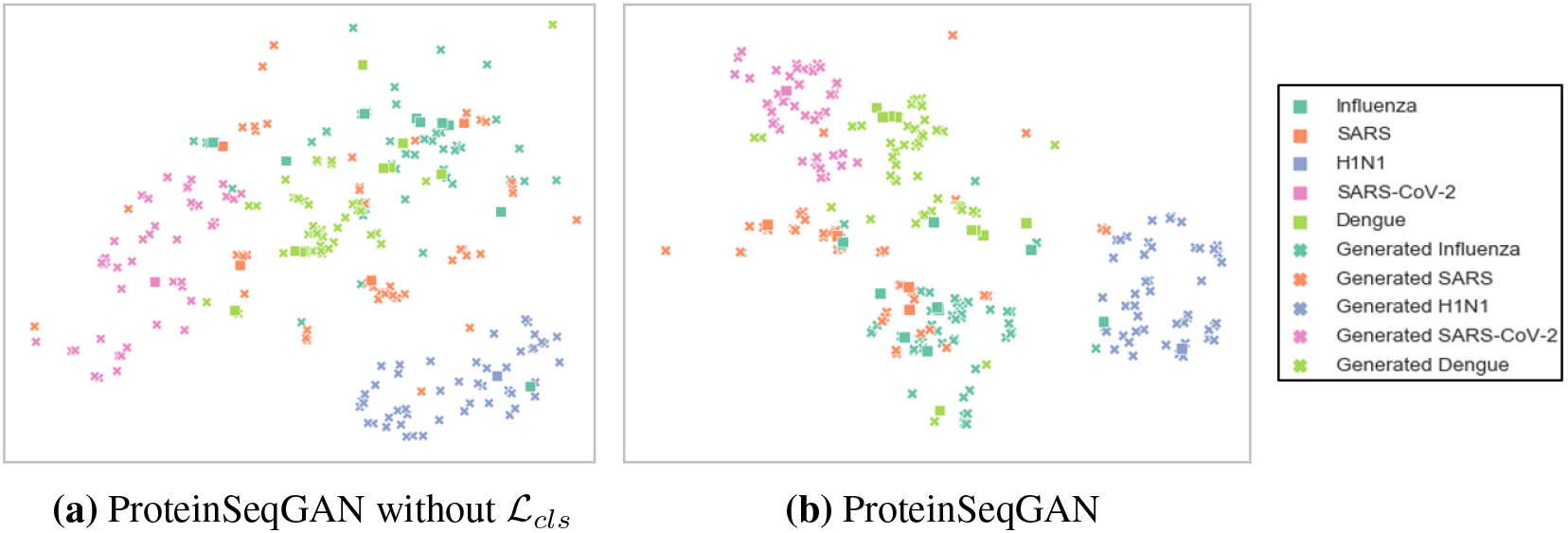
t-SNE visualization [53] of graphically features *f* extracted from real protein sequences in the dataset, along with **(a)** protein sequences generated by ProteinSeqGAN without 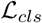, and **(b)** protein sequences generated by ProteinSeqGAN.

### 4.6 Mutation prediction via Sequence Alignment

We present the prediction of virus mutations via the sequence alignment [52] between the generated sequence and its closest related sequence in the dataset, as shown in Table 3, where “-” represents unaligned fragments. By training on the multi-class dataset, our model learns to generated antigen sequences mutated by deleting, inserting, and substituting, which are largely similar to the actual antigens from the corresponding viral species. It is indicated that the generated sequences are biologically relevant to viral species, which can be direct and reliable predictions for virus mutations.

**Table 3:**
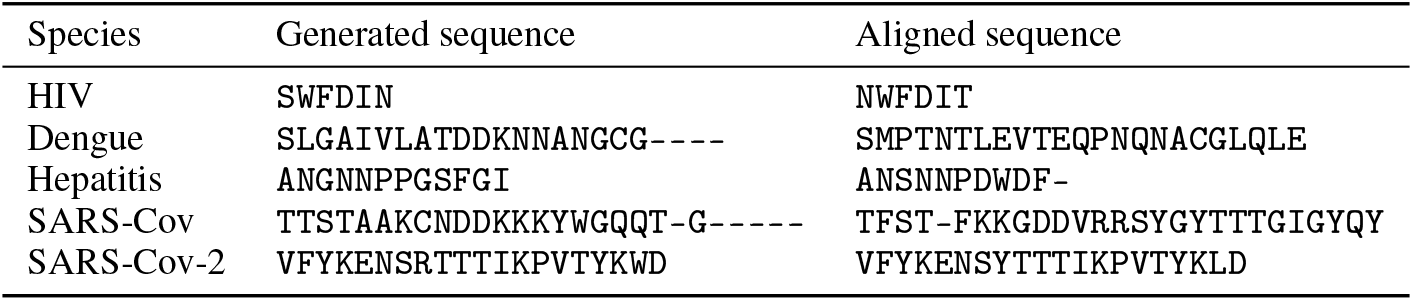
Sequence alignment for mutation prediction

## 5 Conclusion

We have investigated the virus mutation predicted via the antigen segments generation. ProteinSeqGAN, a multi-class protein sequence generative model, has been developed, which is composed of the conditional generator and the GProAE. By graphically featurizing the protein sequences and leveraging the encoder-decoder architecture, the GProAE evaluates the goodness of protein sequences and predict the corresponding viral species. Experiments demonstrate that ProteinSeqGAN can generate valid protein sequences from both bioinformatics and statistics perspectives, and such generated sequences can be direct and reliable predictions for virus mutations.

## Broader Impact

Our ProteinSeqGAN model learns to generate valid viral epitope protein sequences, which can efficiently predict the virus mutations. Through this model, we are also able to predict the possible mutation and evolution of SARS-CoV-2, which is a problem of great interest to humanity [1]. The model can help understand the viral antigen proteins, and also disentangle the connectivity between different virus species.

The epitope mutation generated by our model, which is closely related to the immune reaction [11, 12], is likely to accelerate the design of antibodies [5]. Additionally, by analyzing the generated antigen sequences, researchers can predict the possible mutations which lead to high infectivity and severity. Such a prediction can help researchers and institutions, like the World Health Organization (WHO), prepare beforehand in case a pandemic-like situation arises due to viral mutations [54]. However, since the sequences generated by the model are epitopes [47, 48], this information may not represent the mutations occurring else where in the protein. These mutations can have an allosteric effect on the binding of viral epitope and the antibody. In such circumstances, the prediction generated by ProteinSeqGAN may not be sufficient for predicting the effect of viral mutations.

Implementing the ProteinSeqGAN in generating the whole viral protein sequences is a promising extension, which may be more effective and comprehensive in predicting whole genome viral mutations. Besides, the framework can be applied to not only viral antigen generation but also predicting other sequences, like RNA or DNA chains [55].

## A.1 Amino Acid Feature Matrix

The feature matrix developed for the GProAE model contains various biochemistry and bioinformatics characteristics for each amino acid, which are built on features from RDKit [56] and the data from the AAindex server [57]. Detailed description about all the 38 features are listed below:

1. Hydrophobicity: the indicator of the amino acids whose side chains are reluctant to reside in aqueous environments Hydrophobicity is an important indicator on how different amino acids will interact and is important for protein stabilizing and folding.
2. Membrane buried preference parameter: quantitates the preference of an amino acid to occur in the lipid bilayer of the cells, thereby indicating its preference to occur in the trans-membrane proteins.
3. Average flexibility index: indicates the stereo-chemical flexibility of the amino acids. Higher flexibility index means that an amino acid can “bend” more comparatively.
4. Solvation free energy: the free energy associated with the solvation of the amino acid with water.
5. Number of hydrogen bond donors: corresponds to the number of hydrogen bonds that can be formed by the amino acid by donating a hydrogen atom.
6. Normalized Van der Waals volume: associated with the molecular size of amino acid. Van der Waals volume represents the size of the constituent atoms.
7. Polarity: the quantification of the net polarity of the amino acid associated with its functional group.
8. Ratio of buried and accessible molar fractions: the ratio of amino acids that are buried in the core of the protein and are inaccessible by water molecules to the accessible amino acids which can for interactions with the water.
9. Distance between c-a and centroid of sidechain: the quantification of the size of amino acid. It represents the distance between the alpha carbon atom and the centroid of the attached cide chain.
10. Normalized frequency of *α* helix: the frequency of an amino acid to occur as part of an *α* helix.
11. Normalized frequency of *β* sheet: the frequency of an amino acid to occur as part of a *β* sheet.
12. Side chain orientation: the possible stereo-chemical orientations that the side chain of an amino acid can take.
13. Accessible surface area: the solvent accessible surface area. It is associated with the portion of the amino acid that can interact with the solvent molecules.
14. Loss of hyrdopathy by helix formation: the change in hydrophobicity of amino acid associated with its occurrence in an alpha helix.
15. Activation Gibbs energy unfolding: the free energy that must be supplied to ensure that the given amino acid comes of the secondary structure it has formed, thereby causing unfolding of the protein.
16. Side chain contribution to stability: a measurement of the impact the amino acid has on the stabilization of the protein molecule.
17. Mean volume of residues buried: the volume of the amino acid that becomes solvent inaccessible when it is part of a secondary structure.
18. pKx: pKa or pKb associated with the functional group on the side chain of the amino acid.
19. Aromatic/stacking: indicates whether an amino acid is aromatic or not and whether it can for stacking interactions.
20. Carbon atoms: the number of Carbon atoms in an amino acid.
21. Nitrogen atoms: the number of Nitrogen atoms in an amino acid.
22. Sulphur atoms: the number of Sulphur atoms in an amino acid.
23. Oxygen atoms: the number of Oxygen atoms in an amino acid.
24. Hydrogen atoms: the number of Hydrogen atoms in an amino acid.
25. Rings: the number of rings in an amino acid if present.
26. Ring atoms: the number of atoms presenting in the ring.
27. *sp*^2^ Hybridization: the number of atoms in an amino acid that have *sp*^2^ hybridization.
28. *sp*^3^ Hybridization: the number of atoms in an amino acid that have *sp*^3^ hybridization are indicated by this feature
29. Degree-1 atoms: the number of atoms with one directly bonded neighbours.
30. Degree-2 atoms: the number of atoms with two directly bonded neighbours.
31. Degree-3 atoms: the number of atoms with three directly bonded neighbours.
32. Single bond: the number of single bonded atoms in an amino acid.
33. Double bond: the number of double bonded atoms in an amino acid.
34. Implicit Valency-0: the number of Hydrogen that can bond to an atom in an amino acid. Here, the number of atoms to which 0 Hydrogen atoms can bond.
35. Implicit Valency-1: the number of atoms to which only one 1 Hydrogen atom can bond.
36. Implicit Valency-2: the number of atoms to which only one 2 Hydrogen atom can bond.
37. Implicit Valency-3: the number of atoms present in an amino acid to which only one 3 Hydrogen atom can bond.
38. Mass: the mass of the amino acid.

## A.2 Dataset

The dataset is composed of the CATNAP [47], the VirusNet [48] and the recent SARS-CoV-2 virus sequence [49]. In total, 1934 viral antigen sequences from over 16 different virus species are collected. The sequences from CATNAP dataset are the viral antigen amino acids, which are within 7 angstroms (Å) of the antibody, and in VirusNet, viral antigens are restricted within 5Å from the antibodies. Shown in Table 4 are some antigen amino acid sequences in the dataset. These antigens identifies different viruses, and are closely related to the immunity reaction. Therefore, prediction of mutations in antigens are of great importance for antibody design.

**Table 4:** Antigen sequences in the dataset

| Species | Sequences | Species | Sequences |
| --- | --- | --- | --- |
| Influenza | RVAGIENTIDGWLKTQAIDINLNIE | H1N1 | PTSSWPNEKGSSYPKSKSYVNQNET |
| Influenza | PPEQWEGMIDGRHEGTGQAALNAER | SARS-CoV | TFSTFKKGDDVRRSYGYTTTGIGYQY |
| Dengue | KAETQHGKIE | SARS-CoV | HHQSPTVDGWDKTQIDTVNI |
| Dengue | IDKEMAETQKKVNYTLHWFRKH | SARS-CoV-2 | VFYKENSYTTTIKPVTYKLD |

## A.3 Detailed Criteria for Statistics Validation

After generating the natural vector *v_N_* for each actual protein sequence from the dataset, two criteria are introduced to validate an arbitrary amino acid sequence A [46]. First, if *v_N_* calculated from A is the same as one of the existing sequence in the dataset, then A is valid only if it is exactly the same as the corresponding sequence. Second, when the first criterion is not satisfied, a convex hull is created by the bounds of *v_N_* of actual protein sequences. For each amino acid “k”, all the corresponding entries in *v_N_*, namely *n_k_*, *μ_k_*, and 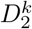, are collected to build a three dimensional convex hull. In total, twenty convex hulls corresponding to all the twenty standard amino acids are built. If the natural vector of an arbitrary amino acid sequence falls within all the convex hulls, it is said to be a valid sequence. Shown in Fig. 5 is the convex hull built on *v_N_* obtained from our training dataset.

**Figure 5:**
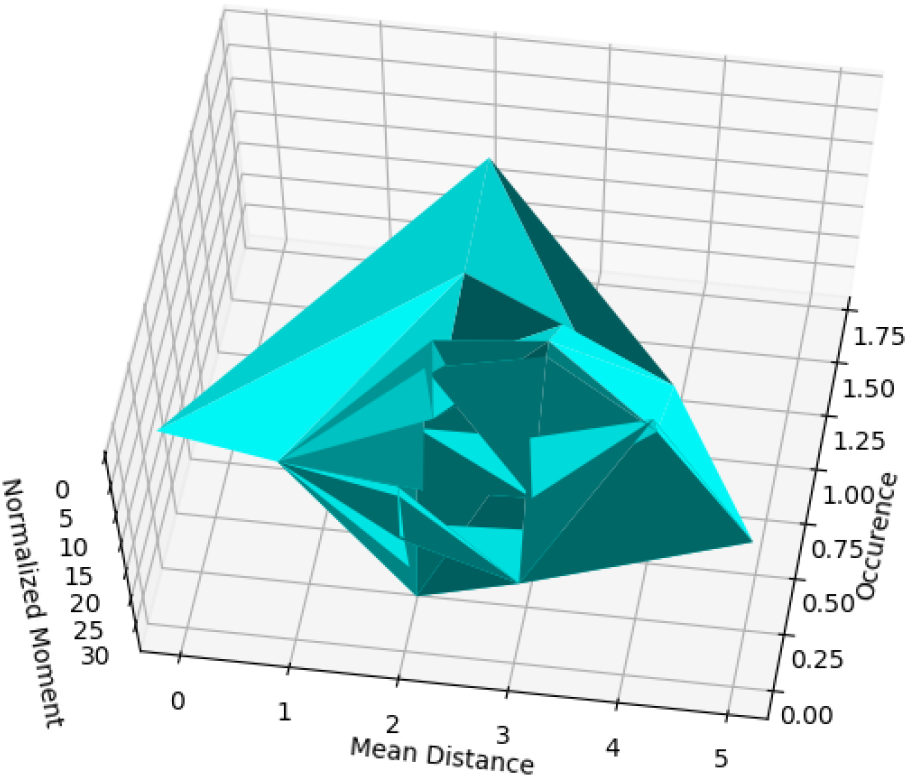
Convex hull built on *v_N_* obtained from our training dataset [47, 48], where the three axes: occurrence, mean distance, normalized moment represent *n_k_, μ_k_*, and 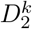 respectively.

## A.4 Graphical Features of Generated Proteins

We also investigate the graphical features extracted from valid sequences generated by ProteinSeqGAN along with actual proteins from all the 16 different viral species. T-SNE [53] is leveraged to map the high dimensional feature vectors to 2D domain. Fig. 6 illustrates that our ProteinSeqGAN can generate various protein sequences which are closely related to the viral species.

**Figure 6:**
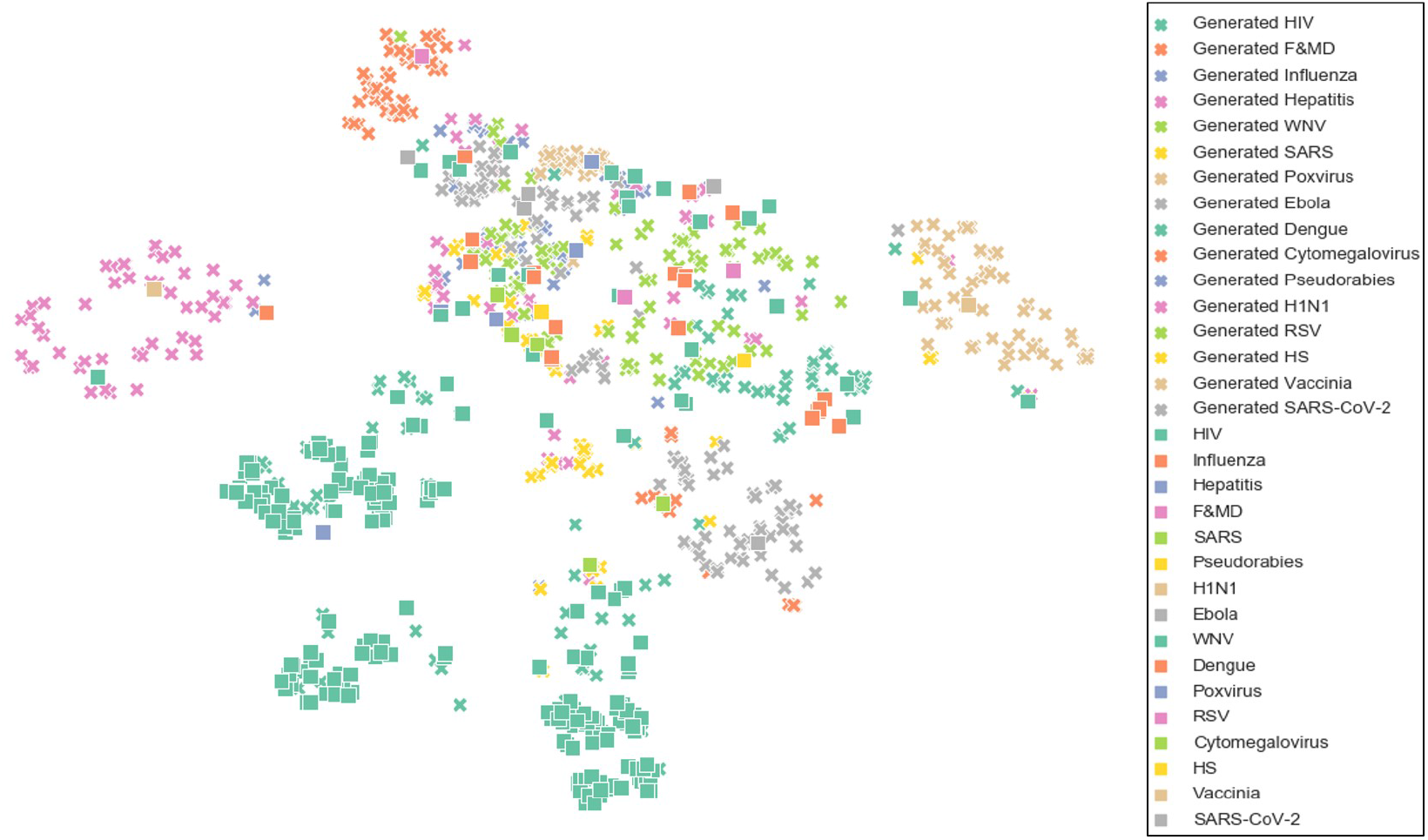
T-SNE visualization [53] of graphically features *f* extracted from protein sequences generated by ProteinSeqGAN, along with real protein sequences from all the viral species in the dataset.

